# Alignment-Free k-mer Analysis Reveals Age-Associated k-mer Signals in Canine Buccal DNA and Cell-Free DNA

**DOI:** 10.64898/2026.10.24.761006

**Authors:** Sonya Hu, Giorgia Del Vecchio, Maciej Długosz, Marek Kokot, Sebastian Deorowicz, Matteo Pellegrini

## Abstract

Aging is associated with progressive molecular and cellular changes that alter chromatin organization, epigenetic regulation, and microbial community structure. While DNA methylation-based biomarkers have demonstrated age-prediction performance, the potential of alignment-free sequence analysis for detecting age-associated signals remains unexplored. We performed a k-mer-based analysis of whole-genome sequencing datasets from two canine cohorts: 1,187 buccal DNA samples and 100 cell-free DNA (cfDNA) samples. Age-associated k-mers were identified through correlation analysis, and sequence motifs and nucleotide composition patterns were characterized across both sample types. Ridge regression models were used to predict age from k-mer frequency profiles, and synthetic datasets were generated under biological compositional assumptions to evaluate potential drivers of the observed signals. Age-associated k-mer signals were identified in both buccal DNA and cfDNA, with substantially stronger patterns observed in buccal samples. Buccal-derived k-mers exhibited more pronounced motif patterns, stronger GC and AT enrichment trends, and improved age-prediction performance relative to cfDNA. Nucleotide enrichment trends differed between sample types, suggesting that distinct biological processes may contribute to age-associated sequence composition changes in buccal DNA and cfDNA. Simulation analyses successfully reproduced major enrichment trends observed empirically, and age-associated signals were largely preserved following restriction to read-start sequence composition. These findings demonstrate that alignment-free k-mer analysis can detect age-associated sequence composition signals in canine DNA and highlight its potential as a computationally efficient complementary approach for aging biomarker discovery.

## 1. Introduction

Aging is a complex biological process characterized by the progressive accumulation of molecular and cellular damage over time. The hallmarks of aging framework proposes twelve interconnected biological processes, including genomic instability, telomere attrition, epigenetic alterations, loss of proteostasis, mitochondrial dysfunction, cellular senescence, chronic inflammation, and dysbiosis, among others, that collectively contribute to age-related functional decline [1, 2]. Together, these processes drive molecular changes that broadly disrupt genome maintenance, cellular function, and tissue homeostasis. Accurate biomarkers of biological aging are useful because chronological age alone does not fully capture inter-individual variation in aging trajectories and provides limited insight into the underlying molecular and cellular state of an organism. Such biomarkers would enable more precise assessment of biological age, facilitate the monitoring of aging interventions, and improve prediction of age-related disease risk. DNA methylation-based epigenetic clocks have emerged as some of the most successful aging biomarkers, demonstrating that molecular signatures can accurately estimate age across diverse tissues and species [3].

Dogs are recognized as a valuable model for aging research due to their shared environment with humans and their susceptibility to many age-related diseases [5], and epigenetic clocks have similarly been developed in this species using DNA methylation patterns [4]. However, whether age-associated molecular changes are also reflected in broader sequence-derived features of whole-genome sequencing data remains largely unexplored, motivating investigation of canine aging using alignment-free sequence analysis approaches.

Cell-free DNA (cfDNA) consists of fragmented DNA molecules released into circulation primarily through apoptosis, necrosis, and cellular turnover. Because cfDNA originates from multiple tissues throughout the body, its composition and fragmentation patterns provide insight into physiological and pathological processes and have emerged as informative biomarkers in diverse clinical applications [6, 9]. Previous studies have demonstrated that age-associated changes in cfDNA characteristics can reflect aspects of biological aging and age-related physiological decline [8].

Another source of DNA that is commonly obtained are buccal samples containing DNA from both host epithelial and immune cells and the oral microbiome, both of which undergo age-associated changes including somatic alterations in host cells and shifts in microbial community composition over time [13–15]. These distinct biological origins suggest that age-associated genomic patterns may differ between cfDNA and buccal sequencing datasets.

Alignment-free methods have become increasingly important in genomics due to their computational efficiency and ability to characterize sequence composition independent of reference genome alignment [7]. Among these approaches, k-mer based methods represent biological sequences through the frequencies of short nucleotide substrings and have been widely applied in genome comparison, metagenomics, and sequence classification [12].

Although aging biomarkers based on DNA methylation have been extensively studied, it remains unclear whether age-associated molecular changes are reflected in broader sequence-derived features of whole-genome sequencing data. Furthermore, the extent to which such patterns vary across biologically distinct sample types, including cfDNA and buccal DNA, is not well understood. In this study, we investigated whether age-associated sequence patterns are detected in canine cfDNA and buccal whole-genome sequencing datasets using an alignment-free k-mer framework. We then evaluated the extent to which these patterns differed between sample types and assessed their utility for age prediction and biological interpretation.

## 2. Methods

### 2.1 Dataset

Whole-genome sequencing (WGS) data from domestic dogs (*Canis lupus familiaris*) were obtained from two independent datasets. The first dataset consists of 1,187 buccal DNA samples from NCBI BioProject PRJNA1525050. The second dataset consists of 100 plasma-derived cell-free DNA samples obtained from the NCBI BioProject PRJNA1373368. Chronological age information was obtained from associated sample metadata and used as the target variable for downstream analysis. Publicly available sequencing data were downloaded from the NCBI Sequence Read Archive (SRA) using SRA Toolkit. All sequencing reads were obtained in FASTQ format and processed using the same preprocessing workflow.

### 2.2 Sequence Preprocessing

To analyze the effects of read preprocessing on k-mer-based age prediction, two approaches were examined: raw sequencing reads and quality-filtered, truncated reads. For the raw-read analysis, FASTQ files were used directly without additional preprocessing. For the truncated-read analysis, FASTQ files were processed using fastp with a minimum Phred quality score of 20 and a minimum retained read length of 7 bp. Adapter trimming was disabled. Following quality filtering, reads were truncated to the first 7 nucleotides, generating fixed-length FASTQ sequences for analysis.

### 2.3 K-mer Frequency Analysis with MKMC

MKMC v0.21.0 utilizes KMC to count k-mers in each input sample [20]. It combines individual KMC databases into a single k-mer matrix, in which the rows correspond to unique k-mers and columns correspond to samples. The source code and documentation for MKMC are available through its GitHub repository: https://github.com/refresh-bio/MKMC. Matrix entries represent the abundance of each k-mer within each sample.

To analyze the effects of k-mer length on age-associated sequence patterns, k-mer sizes of 6, 7, and 8 were tested using raw buccal DNA sequencing reads. Spearman correlation analysis was performed between k-mer frequencies and chronological age, with Bonferroni correction applied for multiple testing.

For subsequent analyses, MKMC was applied to canine whole-genome sequencing FASTQ files derived from buccal DNA and plasma cfDNA samples using a k-mer size of 7 (-k 7) and matrix output ( -o matrix ). K-mer frequencies were then normalized using MKMC frequency normalization method ( -n freq ) before Spearman correlation analysis (--cor spearman) with chronological age values provided in a phenotype file. The resulting k-mer frequency matrix and correlation statistics were used for age-prediction analyses.

### 2.4 Motif Analysis

To characterize sequence patterns associated with age, motif analysis was performed on k-mers exhibiting the strongest Spearman correlations with chronological age. For truncated datasets, the 100 most positively correlated and 100 most negatively correlated k-mers were used as input sequences for Weblogo to visualize conserved nucleotide patterns. For raw datasets, de novo motif discovery was performed on the same k-mer subsets using the MEME Suite. Motif discovery was performed using MEME in Classic mode with the zero-or-one occurrence per sequence (ZOOPS) model, searching both DNA strands with motif widths restricted to 7-10 nucleotides. Statistically significant motifs (E-value < 0.05) were kept for interpretation.

### 2.5 Age Prediction

Age prediction was performed using Ridge Regression using scikit-learn v1.6.1 within Python 3.10.12, with k-mer frequency matrices generated by MKMC used as input features and chronological age as the target variable. All k-mer features from the normalized frequency matrix were included in model training without prior correlation-based feature selection. To address the high-dimensional feature space, Ridge Regression with L2 regularization was applied. The regularization parameter, α, was optimized using RidgeCV over a logarithmically spaced grid of 30 values ranging from 10^-4^ to 10^4^, with optimal α selected independently within each training fold. Spearman correlation-based ranking of age-associated k-mers was performed separately for exploratory analysis and was not used to select predictors for the age-prediction models.

Model performance was evaluated using leave-one-out cross-validation (LOOCV) for both buccal DNA (n = 1187) and cfDNA (n=100) datasets to maintain consistent evaluation strategy across datasets. In each iteration, one sample was held out for testing while the remaining samples were used for training. Prediction accuracy was assessed using the coefficient of determination (R²), root mean squared error (RMSE), mean absolute error (MAE), and Spearman rank correlation coefficient (ρ) between predicted and chronological age. Scatter plots of predicted versus chronological age were generated to visualize the relationship between predicted and chronological age and assess systematic prediction bias.

### 2.6 Synthetic Sequence Simulation

Synthetic sequencing datasets were generated using a custom Python pipeline to evaluate whether the observed nucleotide enrichment patterns could be reproduced under defined compositional models. Synthetic source sequences were computationally generated DNA sequences with predefined GC contents and, where applicable, CpG observed/expected ratios. Host-like and bacterial-like sequences were generated at lengths of 2,000,000 bp each while CpG island-like and shore-like sequences were generated at lengths of 500,000 bp each. These synthetic sequences were designed to model predefined nucleotide composition patterns, rather than replicate complete biological genomes. Nucleotides were randomly shuffled to achieve the target GC content and CpG frequencies were subsequently adjusted through nucleotide substitutions. These sequences served as templates from which 50-bp sequencing reads were randomly sampled. Simulations were performed separately for buccal DNA and cfDNA. To approximate the relative sample sizes of the empirical datasets, 300 buccal samples and 90 cfDNA samples were generated. For each simulated sample, 5,000 sequencing reads were randomly sampled from synthetic source sequences representing age-associated compositional changes.

For buccal DNA, synthetic reads were sampled from host-like and bacterial-like source sequences. The host-like sequence was generated with a target GC content of 40% and a CpG observed/expected ratio of 0.10, while the bacterial-like sequence was generated with a target GC content of 65%. Age-associated compositional shifts were modeled by increasing the proportion of bacterial-like reads across age groups (young: 2 years, middle-age: 8 years, old: 14 years). Young samples contained 4,500 host-like and 500 bacterial-like reads, middle-aged samples contained 3,500 host-like and 1,500 bacterial-like reads, and old samples contained 2,500 host-like and 2,500 bacterial-like reads.

For cfDNA, synthetic reads were sampled from CpG island-like and CpG shore-like source sequences. These regions were selected because CpG sites are associated with age-related epigenetic change [3,4, 19], while cfDNA fragmentation patterns reflect nucleosome positioning and chromatin organization [6,9]. Together, these findings provide a biological rationale for investigating whether age-associated differences in the representation of CpG-rich and CpG-poor sequences could influence k-mer composition. CpG island-like sequences were generated with a target GC content of 55% and a CpG observed/expected ratio of 0.80, whereas CpG shore-like sequences were generated with a target GC content of 42% and a CpG observed/expected ratio of 0.30. To evaluate whether MKMC could detect predefined changes in sequence composition, the proportion of CpG island-like reads was systematically decreased across simulated age groups, from 85% in young samples to 55% in middle-aged samples and 25% in old samples. These proportions were selected as predefined simulation parameters rather than experimentally established age-associated changes in canine cfDNA. Therefore, this simulation evaluates MKMC’s sensitivity to controlled compositional shifts rather than validating its performance on real canine cfDNA.

Synthetic FASTQ files were analyzed using the same MKMC-based k-mer workflow applied to the empirical datasets. MKMC was run with k = 7, normalized k-mer frequencies, and Spearman correlation against age. The top 100 positively and negatively correlated k-mers from each synthetic dataset were used to generate sequence logos with Logomaker and compared with the corresponding empirical sequence logos.

### 2.7 Data Availability

The datasets used in this study are available as follows:

**Whole-genome sequencing (WGS) buccal DNA:** NCBI BioProject PRJNA1525050 (https://www.ncbi.nlm.nih.gov/bioproject/PRJNA1525050).

**Plasma-derived cell-free DNA (cfDNA):** NCBI BioProject PRJNA1373368 (https://www.ncbi.nlm.nih.gov/bioproject/PRJNA1373368).

### 2.8 Code Availability

All custom scripts used for motif analysis, statistical analyses, machine learning, and figure generation are publicly available at https://github.com/Hungrycat112/mkmc_analysis. K-mer counting was performed using MKMC, available at https://github.com/refresh-bio/MKMC.

## 3. Results

To investigate whether aging is associated with changes in DNA sequence composition, we analyzed whole-genome sequencing datasets derived from two biologically distinct sample types: cell-free DNA (cfDNA) and buccal samples. Age-associated changes in DNA sequence composition may arise through several mechanisms, including shifts in DNA coverage driven by processes such as telomere shortening, changes in fragmentation patterns, or age-related alterations in microbial DNA composition. Cell-free DNA represents DNA shed from cells into the bloodstream. It is fragmented, nucleosome-bound, and derived from a mixture of cell types. Buccal samples, in contrast, contain host oral epithelial DNA alongside microbial DNA from the oral microbiome, enabling assessment of age-associated sequence composition patterns in a sample type with substantial microbial content. Age-associated changes in sequence composition were assessed by computing k-mers and quantifying their abundance across samples.

### 3.1 Study Design and k-mer Size Selection

Whole-genome sequencing data were obtained from two canine datasets: a cell-free DNA fragmentation study (NCBI BioProject PRJNA1373368; n = 100), and a buccal DNA sequencing dataset (NCBI BioProject PRJNA1525050; n = 1,187). FASTQ reads from both sample types were processed using an alignment-free k-mer analysis pipeline (Figure 1). Reads were analyzed under two preprocessing conditions: raw reads retaining their original read length without preprocessing and truncated reads subjected to quality filtering using fastp (v1.1.0; Phred ≥ 20, minimum retained length of 7 bp, adapter trimming disabled), followed by truncation to the first seven nucleotides. Analysis of raw reads was used to assess age-associated variation in k-mer abundance and sequence composition, whereas analysis restricted to the first seven bases was used to evaluate whether fragmentation-associated or read-start compositional patterns contributed to the observed signals.

**Figure 1.**
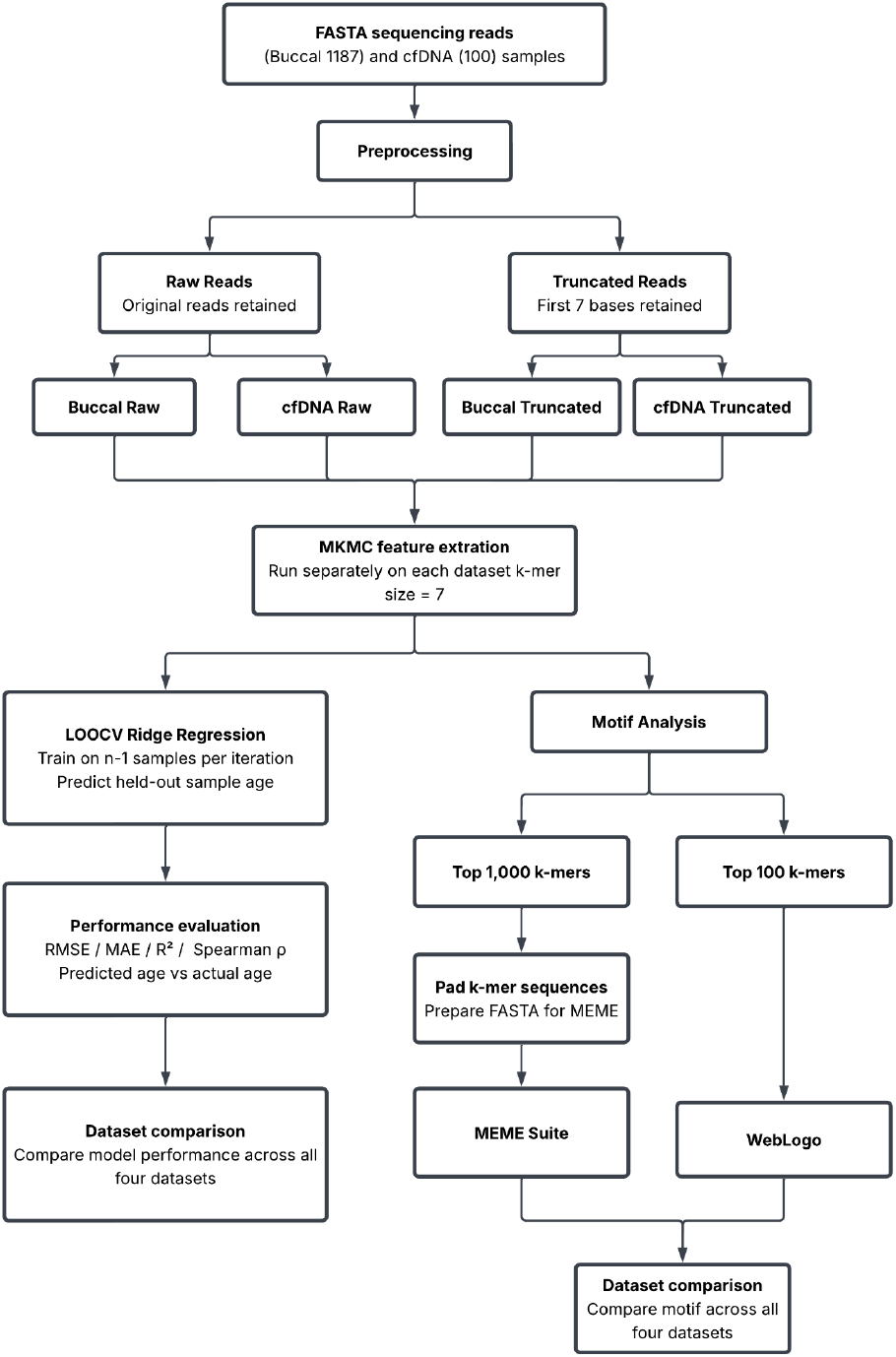
Workflow for k-mer Based Age Prediction and Motif Analysis

An overview of the complete workflow is shown in Figure 1. K-mer frequency matrices were generated using MKMC, which counts k-mers across individual samples. Resulting k-mer frequencies were combined into a single matrix with k-mers as rows, samples as columns, and entries representing per-sample k-mer counts. Spearman correlation analysis and LOOCV ridge regression were then performed as postprocessing steps within the MKMC framework to quantify age associations and evaluate age prediction performance from k-mer frequency profiles. Motif analysis of the top 100 age-associated k-mers was conducted using MEME Suite and WebLogo.

Buccal and cfDNA datasets differed substantially in both sample size and age distribution (Figure 2). Buccal samples were skewed toward younger ages, with a median age of 1.50 years (IQR = 0.58–4.21 years; range = 0.02–17.00 years), whereas cfDNA samples were concentrated among older individuals, with a median age of 10.00 years (IQR = 8.00–12.00 years; range = 3.00–15.00 years). The buccal and cfDNA datasets contained 1,187 and 100 samples, respectively. These differences in sample size and age distribution should be considered when comparing age-associated patterns between datasets.

**Figure 2.**
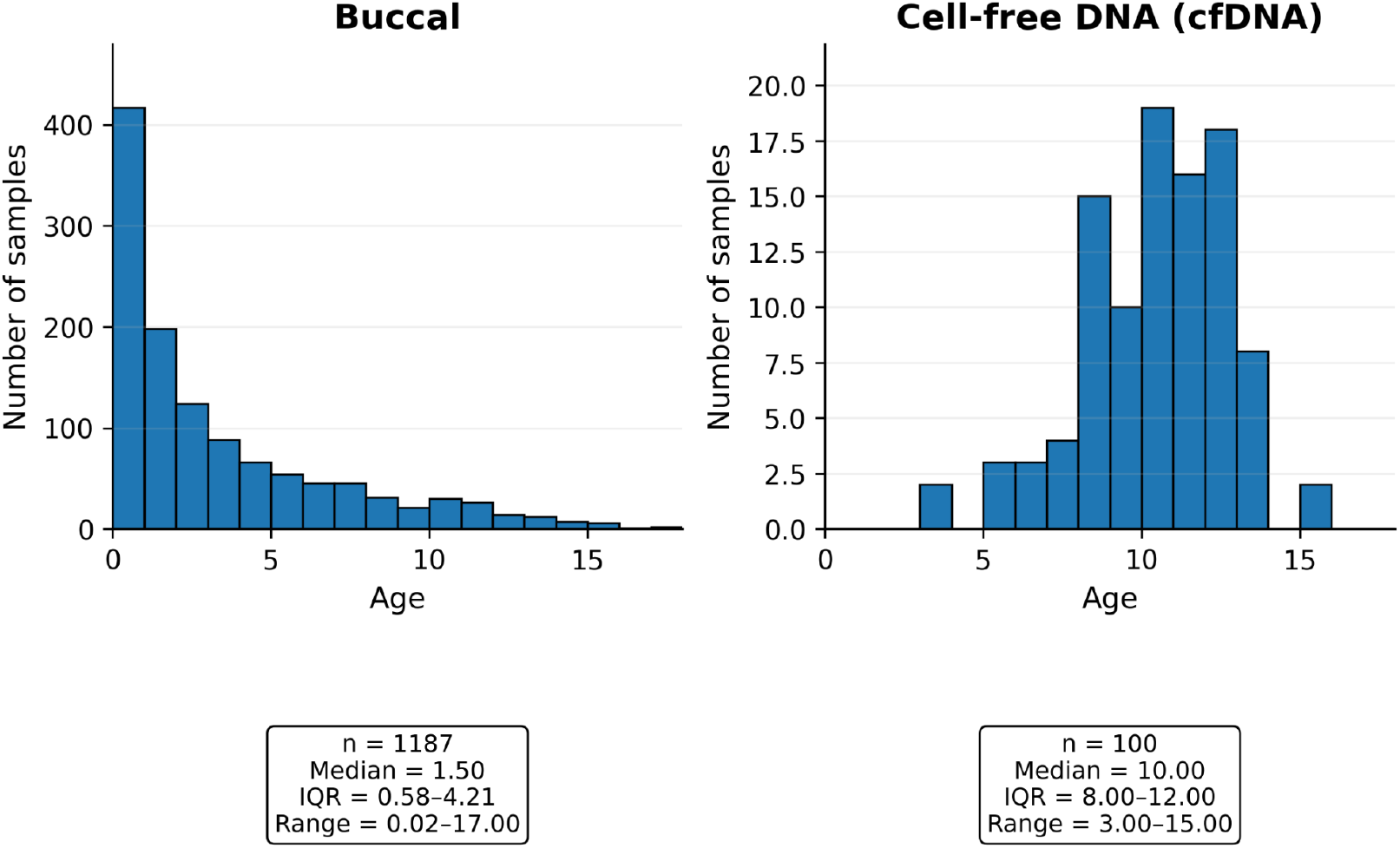
Age distributions of canine buccal and cell-free DNA (cfDNA) cohorts

To assess appropriate k-mer size selection, Spearman correlation analyses were performed on raw buccal datasets using k-mer sizes of 6 and 8 (Supplementary Figure 1). The number of Bonferroni-corrected significant k-mers increased with k-mer size (k = 6: 1,763; k = 7: 7,045; k = 8: 28,583), consistent with the larger total k-mer space at higher k values. Although k = 6 exhibited a similar overall correlation distribution, its substantially smaller k-mer space (n = 2,080 vs. n = 8,192 for k = 7) yielded fewer significant age-associated k-mers. Together, these results indicate that the major age-associated sequence composition patterns were preserved across k-mer sizes while supporting the use of k = 7 for subsequent analyses. Relative to k = 6, k = 7 provides greater k-mer diversity while retaining the overall correlation structure observed across age-associated k-mers.

### 3.2 Age-Associated k-mer Correlation Patterns Across Raw cfDNA and Buccal Datasets

To analyze age-association sequence composition patterns in full-length sequencing reads, Spearman correlation analysis was performed on raw cfDNA and buccal datasets (Figure 3). Buccal samples displayed a strong bimodal correlation distribution, with substantial enrichment of both positively and negatively age-associated k-mers. In contrast, cfDNA samples had a narrower correlation structure with reduced correlation magnitude overall.

**Figure 3.**
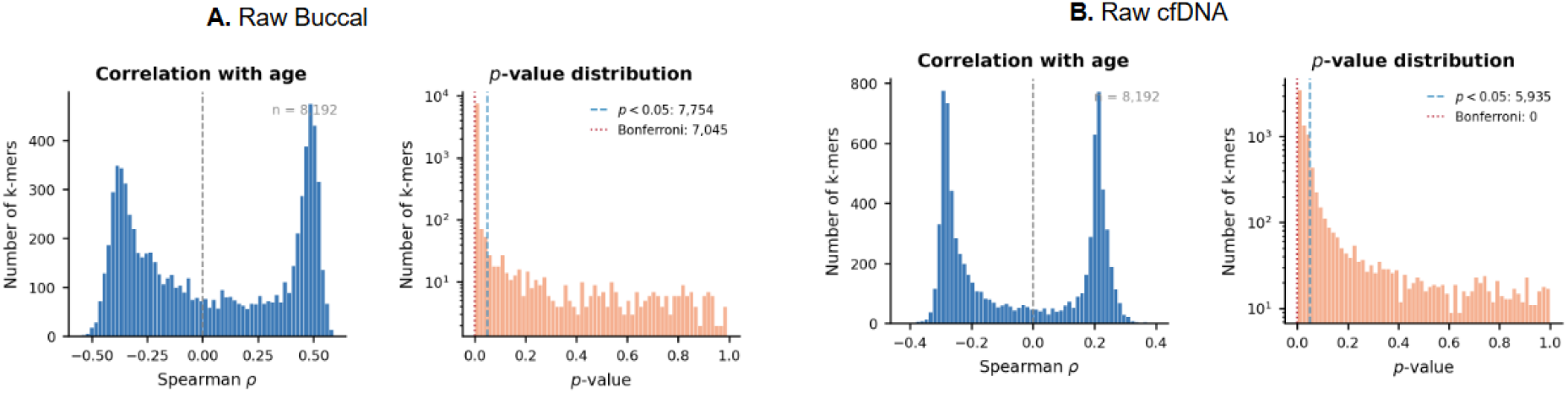
Distribution of age-associated k-mer correlations and p-values across raw cfDNA and buccal datasets

The raw buccal dataset contained 7,045 Bonferroni-significant k-mers, whereas no Bonferroni-significant k-mers were identified in the raw cfDNA dataset despite 5,935 k-mers passing the nominal significance threshold of p < 0.05 (Figure 3). These findings indicate more significant age-associated sequence composition signals in buccal-derived DNA relative to cfDNA-derived DNA.

To evaluate whether these correlation patterns translated into age predictive models, leave-one-out cross-validation (LOOCV) ridge regression was performed using k-mer frequency profiles (Figure 4). Age prediction performance was substantially stronger in raw buccal samples (Spearman ρ = 0.544, R² = 0.270, RMSE = 2.96) compared with raw cfDNA samples (Spearman ρ = 0.215, R² = 0.037, RMSE = 2.27). Although cfDNA samples showed statistically significant associations, predictive performance remained weak relative to buccal datasets.

**Figure 4.**
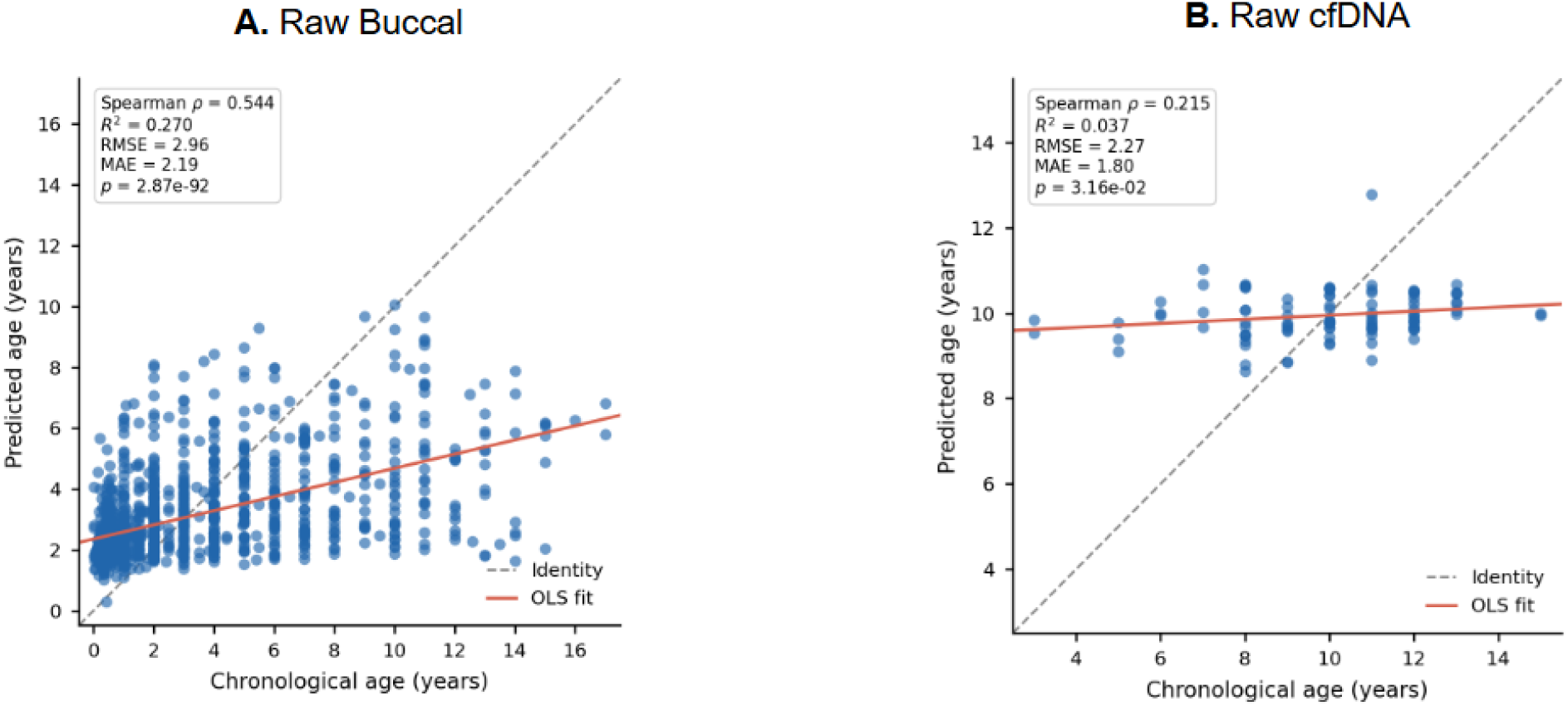
LOOCV ridge regression age prediction in raw cfDNA and buccal datasets

Together, these results demonstrate that age-associated k-mer signals are substantially stronger in buccal than cfDNA-derived DNA, with buccal samples exhibiting greater correlation magnitude, more Bonferroni-significant k-mers, and overall better age prediction performance.

### 3.3 Age-Associated k-mer Correlation Patterns Across Truncated cfDNA and Buccal Datasets

To examine whether age-associated sequence patterns are observed at the start of reads due to changes in fragmentation preferences, Spearman correlation analysis was performed on truncated datasets containing only the first seven bases of each sequencing read (Figure 5). Similar to the raw datasets, truncated buccal samples exhibited a strongly bimodal correlation structure with substantial enrichment of both positively and negatively age-associated k-mers. In contrast, truncated cfDNA samples displayed weaker overall correlation structure and substantially reduced correlation magnitude.

**Figure 5.**
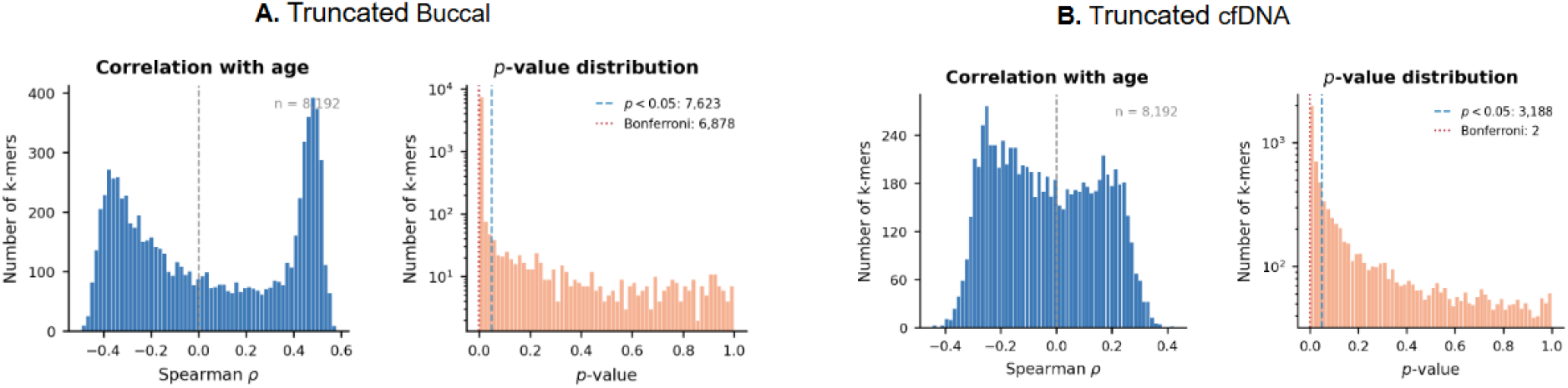
Distribution of age-associated k-mer correlations and p-values across truncated cfDNA and buccal datasets

The truncated buccal dataset contained 6,878 Bonferroni-significant k-mers, whereas only two Bonferroni-significant k-mers were identified in the truncated cfDNA dataset despite 3,188 k-mers passing the nominal significance threshold of p < 0.05 (Figure 5). Correlation distributions and significance patterns were broadly consistent with raw datasets, indicating that the major age-associated sequence composition trends were preserved following truncating.

LOOCV ridge regression on truncated k-mer frequency profiles similarly demonstrated substantially stronger age-predictive performance in buccal samples (ρ = 0.545, R² = 0.267, RMSE = 2.97) relative to cfDNA samples (ρ = 0.210, R² = 0.015, RMSE = 2.30) (Figure 6). Predictive performance in truncated buccal datasets remained highly similar to raw analyses, whereas cfDNA datasets continued to exhibit comparatively weak predictive signals.

**Figure 6.**
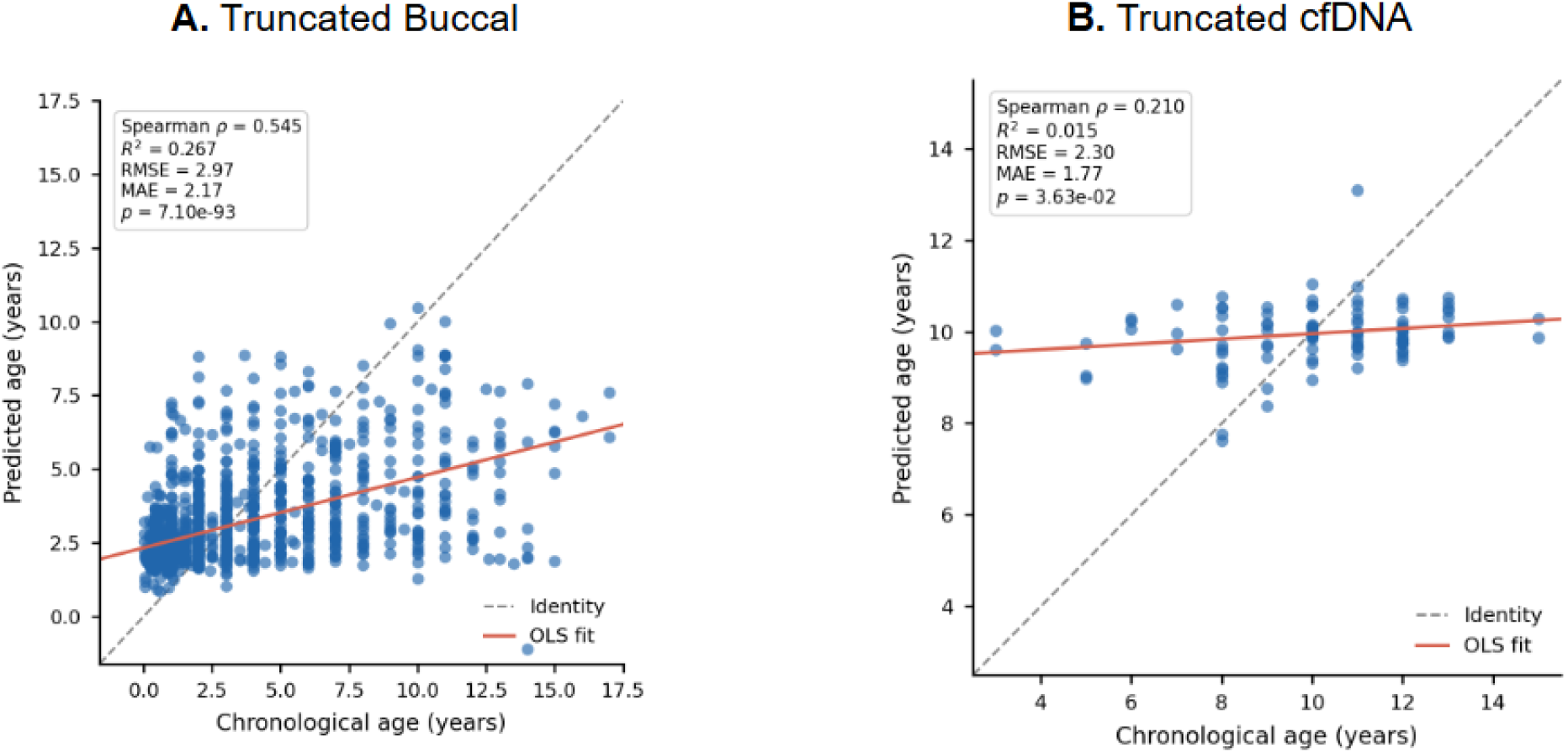
LOOCV ridge regression age prediction in truncated cfDNA and buccal datasets

Together, these findings indicate that age-associated k-mer signals in buccal DNA are preserved when analysis is restricted to read-start positions, suggesting that read-start sequence composition alone is sufficient to capture the age-associated signal present in full-length reads.

Table 1 summarizes LOOCV ridge regression performance across all datasets. Consistent with the results presented in the preceding sections, buccal datasets exhibited substantially stronger age-predictive performance than cfDNA datasets, whereas truncation had minimal impact on predictive accuracy within each sample type. Together, these findings indicate that sample source was the primary determinant of model performance.

**Table 1.** Summary of LOOCV Ridge regression performance across sample types.

| Sample | Spearman $\rho$ | $R^2$ | RMSE | MAE | p-value |
| --- | --- | --- | --- | --- | --- |
| Truncated Buccal | 0.545 | 0.267 | 2.97 | 2.17 | 7.10e-93 |
| Raw Buccal | 0.544 | 0.270 | 2.96 | 2.19 | 2.87e-92 |
| Truncated cfDNA | 0.210 | 0.015 | 2.30 | 1.77 | 3.63e-02 |
| Raw cfDNA | 0.215 | 0.037 | 2.27 | 1.80 | 3.16e-02 |

### 3.4 Motif Analysis and Sequence Composition Trends

MEME motif analysis of the top 100 positively and negatively correlated k-mers from each raw dataset identified statistically significant sequence motifs across all tissue types and correlation directions, with distinct levels of conservation and statistical support between tissues (Figure 7). Buccal samples yielded highly significant motifs in both correlation directions (ρ < 0: E-value = 5.9×10⁻⁵⁶; ρ > 0: E-value = 6.3×10⁻⁴⁵), with well-defined positional conservation across multiple sites, consistent with the GC-enriched motif observed in positively correlated k-mers (Panel B) and mixed nucleotide composition in negatively correlated k-mers (Panel A). By contrast, cfDNA-derived motifs were substantially weaker, with lower positional information content and less significant E-values (ρ < 0: E-value = 2.3×10⁻¹⁴; ρ > 0: E-value = 7.8×10⁻⁵), and motif signal was concentrated at fewer positions, indicating reduced sequence conservation among age-associated k-mers in cfDNA. Collectively, these findings reinforce the nucleotide composition trends observed in analysis of logos and indicate stronger positional motif conservation among buccal-derived age-associated k-mers relative to cfDNA.

**Figure 7:**
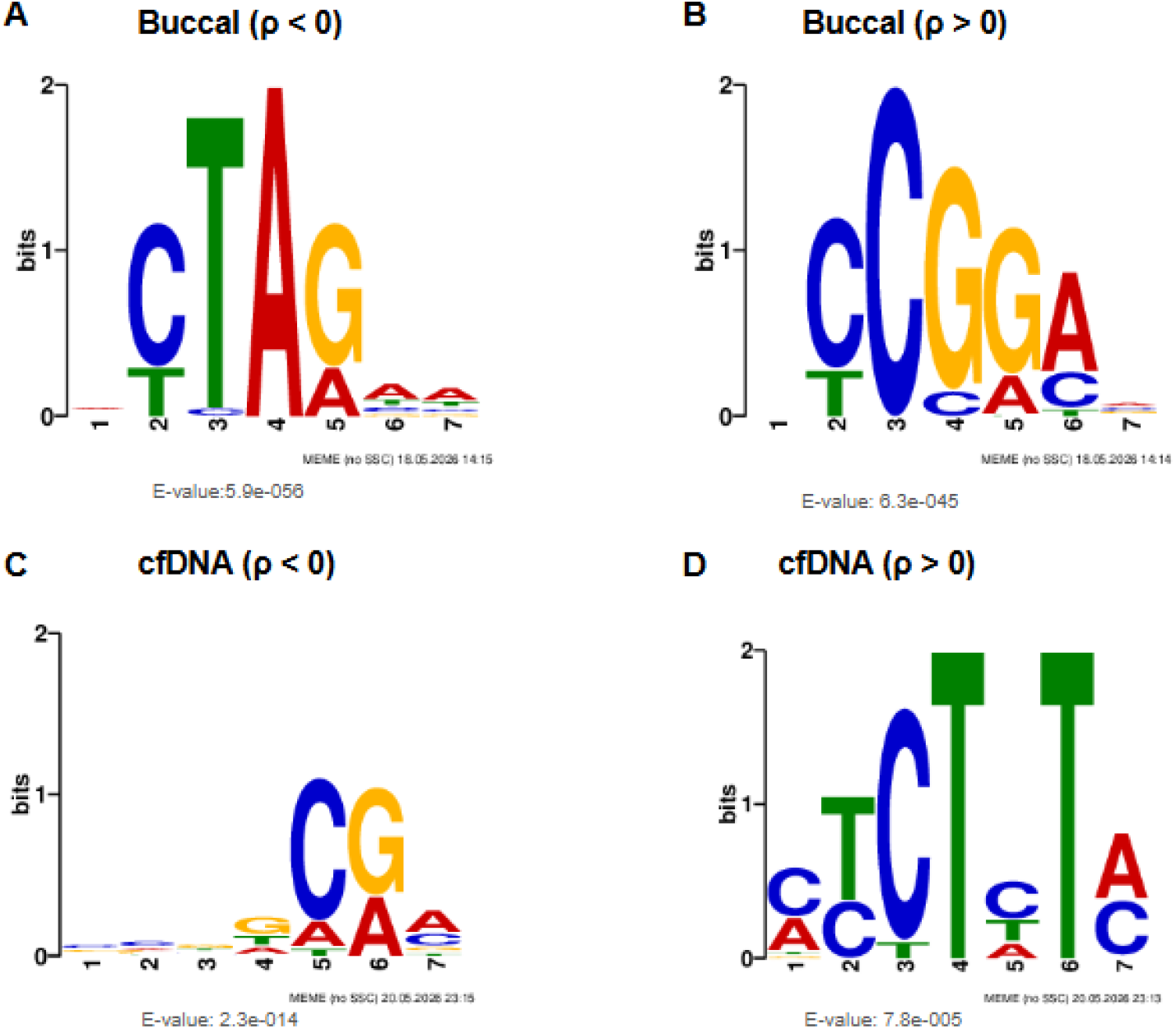
Motifs identified from raw sequencing data using MEME (top 100 k-mers). (A) Buccal samples with negatively correlated k-mers (ρ < 0). (B) Buccal samples with positively correlated k-mers (ρ > 0). (C) cfDNA samples with negatively correlated k-mers (ρ < 0). (D) cfDNA samples with positively correlated k-mers (ρ > 0).

To assess whether motif signals were robust to k-mer set size, MEME analysis was additionally performed on the top 1,000 positively and negatively correlated k-mers from raw datasets (Supplementary Figure 2). Buccal samples continued to yield statistically significant motifs in both correlation directions, with two distinct motifs identified among negatively correlated k-mers (ρ < 0; E-value = 1.9 × 10⁻⁵⁷ and 4.8 × 10⁻⁶) and one among positively correlated k-mers (ρ > 0; E-value = 1.6 × 10⁻⁶¹), consistent with patterns identified in the top 100 k-mer analysis. In contrast, no statistically significant motif was detected among positively correlated cfDNA k-mers under this expanded set. These results suggest that age-associated sequence patterns were more consistently captured by motif analysis in buccal samples than in cfDNA samples.

Sequence analysis of the top 100 positively and negatively correlated k-mers from each truncated dataset revealed distinct nucleotide composition patterns between buccal and cfDNA samples (Figure 8). In buccal samples, positively correlated k-mers (ρ > 0) were enriched for GC content across most positions (Panel C), whereas negatively correlated k-mers (ρ < 0) exhibited AT enrichment with mixed nucleotide composition at several positions (Panel A). In cfDNA samples, negatively correlated k-mers (ρ < 0) were strongly GC-enriched (Panel B), whereas positively correlated k-mers (ρ > 0) were predominantly AT-rich (Panel D). These contrasting compositional profiles indicate that age-associated nucleotide enrichment patterns differ substantially between buccal- and cfDNA-derived k-mers.

**Figure 8:**
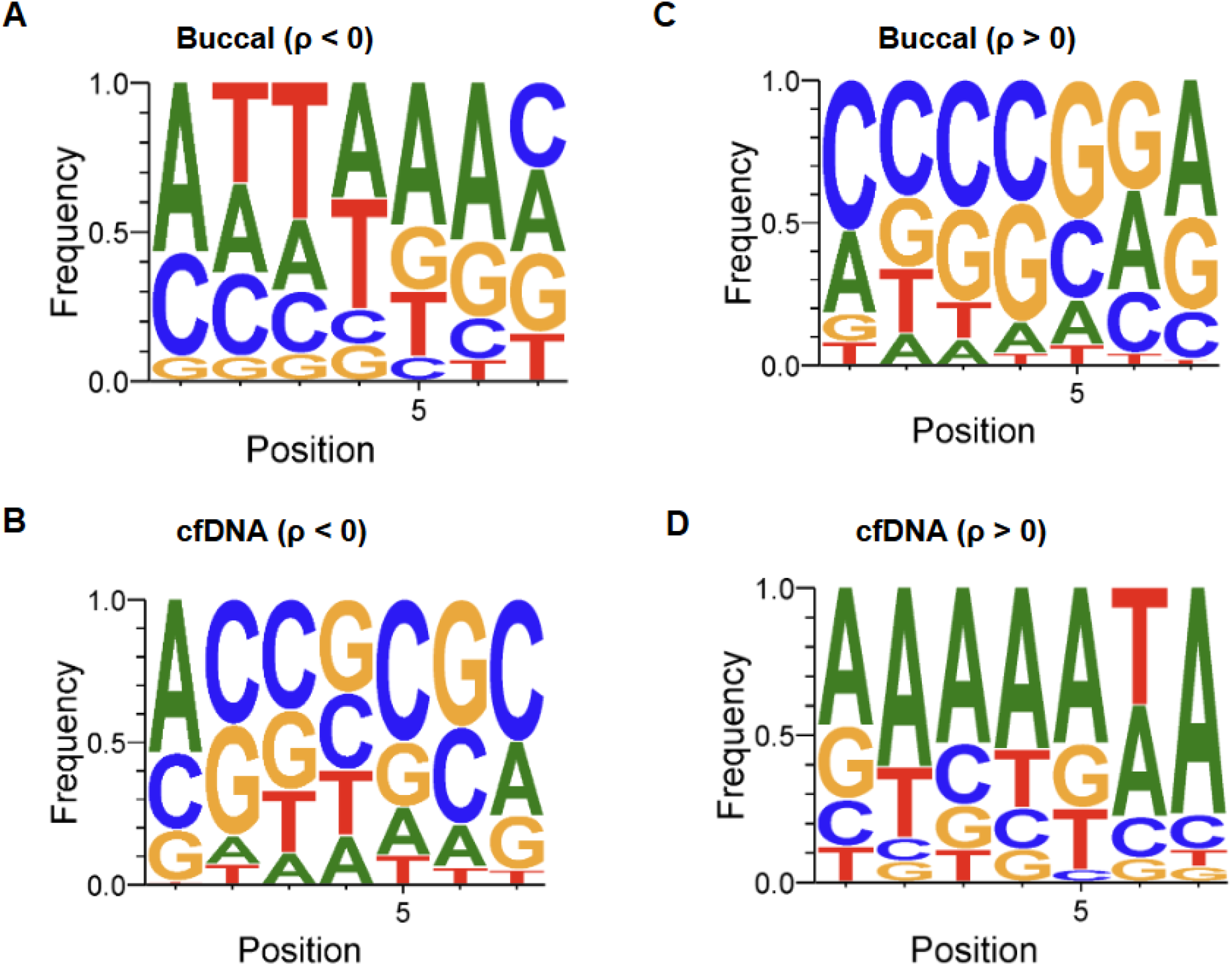
Tissue-specific nucleotide enrichment patterns in age-associated truncated k-mers.

### 3.5 Synthetic Dataset Simulations

To evaluate whether the observed nucleotide enrichment patterns could be reproduced under specific models, synthetic sequencing datasets were generated using a defined simulation pipeline incorporating dataset-specific sequence composition parameters (Figure 9). Sequence logo analysis of synthetic k-mers reproduced the major nucleotide enrichment trends observed in real data across all four tissue-correlation conditions (Figure 10). Specifically, synthetic buccal positively correlated k-mers (Panel E) reproduced the GC-rich motif observed in real buccal data (Panel A), whereas synthetic buccal negatively correlated k-mers (Panel F) reproduced the AT-rich patterns observed in real data (Panel B). Similarly, synthetic cfDNA positively correlated k-mers (Panel G) reflected the AT-rich composition observed in real cfDNA data (Panel C), while synthetic cfDNA negatively correlated k-mers (Panel H) reflected the GC-enriched patterns observed in real cfDNA data (Panel D). The consistency between real and synthetic sequence logos across all sample-type and correlation conditions suggests that the imposed compositional parameters were sufficient to reproduce the major nucleotide enrichment trends observed in the empirical datasets.

**Figure 9:**
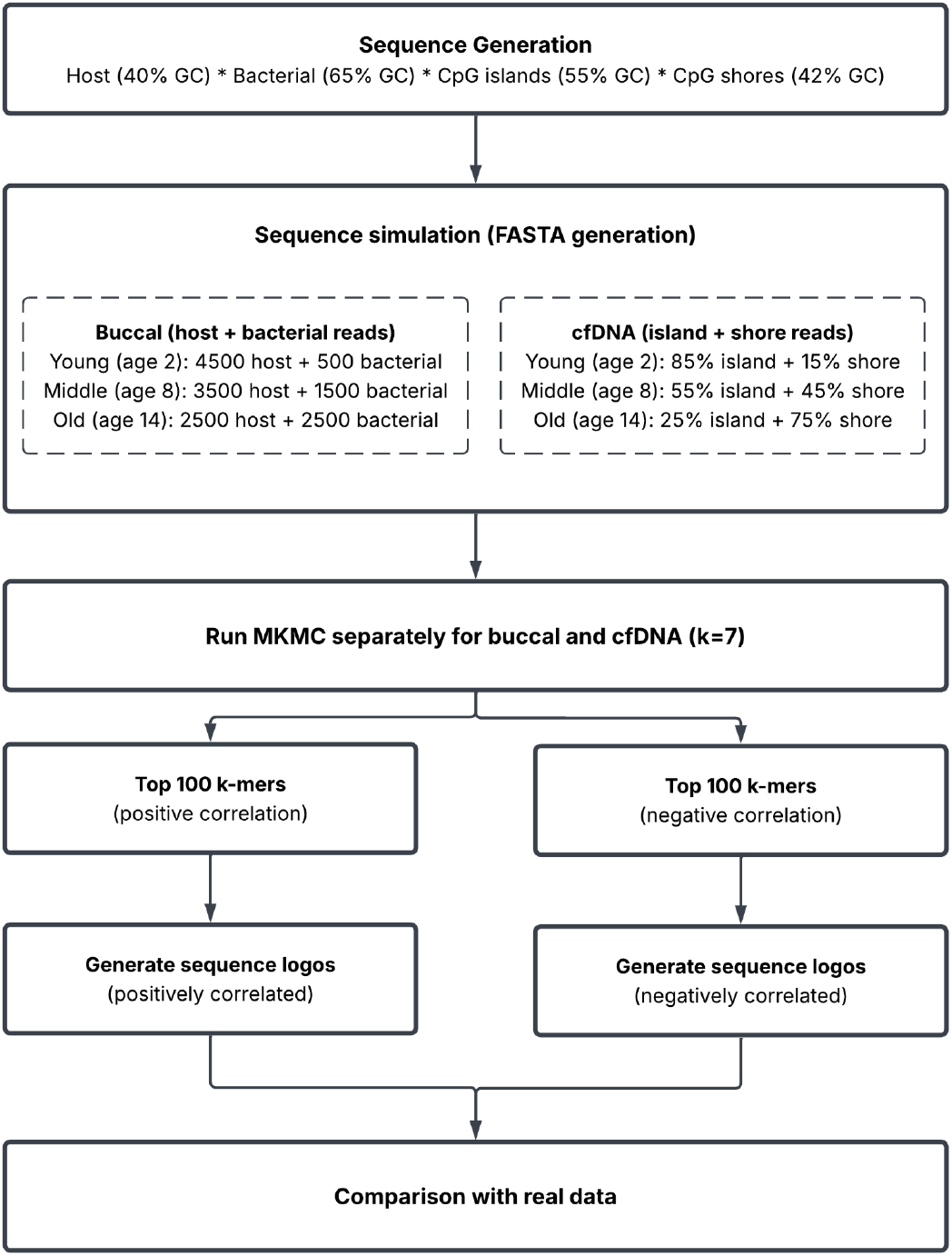
Overview of the synthetic data generation and k-mer analysis pipeline.

**Figure 10:**
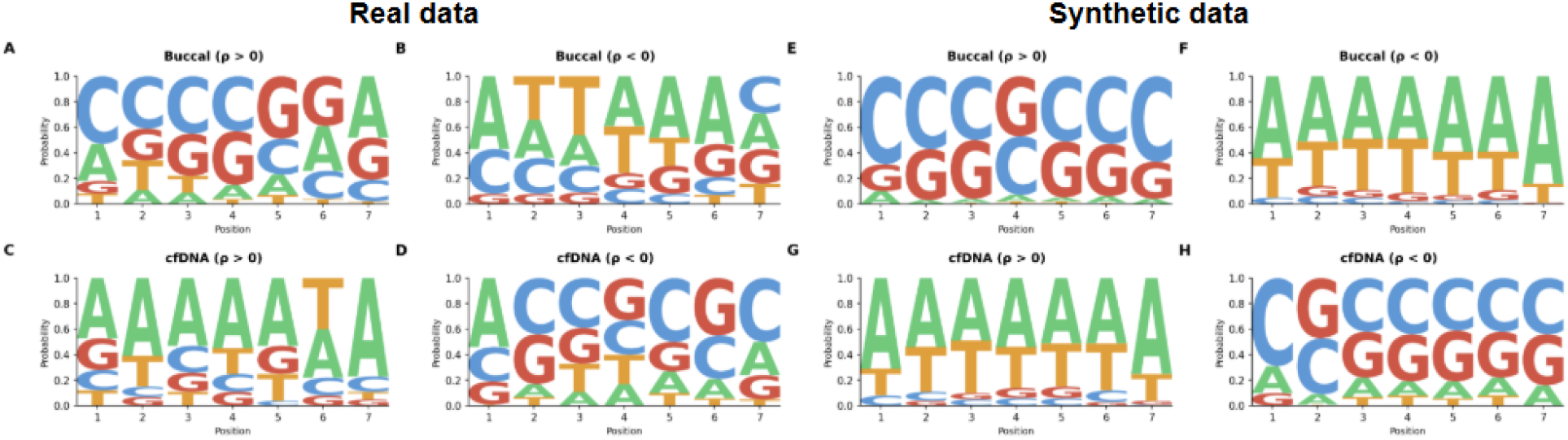
Comparison of sequence logo patterns derived from real and synthetic sequencing data. (A) Real buccal samples with positively correlated k-mers (ρ > 0). (B) Real buccal samples with negatively correlated k-mers (ρ < 0). (C) Real cfDNA samples with positively correlated k-mers (ρ > 0). (D) Real cfDNA samples with negatively correlated k-mers (ρ < 0). (E) Synthetic buccal samples with positively correlated k-mers (ρ > 0). (F) Synthetic buccal samples with negatively correlated k-mers (ρ < 0). (G) Synthetic cfDNA samples with positively correlated k-mers (ρ > 0). (H) Synthetic cfDNA samples with negatively correlated k-mers (ρ < 0).

Overall, these analyses show that age-associated k-mer patterns differ between buccal and cfDNA sequencing datasets, with buccal samples consistently exhibiting stronger correlation structure, greater motif conservation, and improved predictive performance relative to cfDNA. Although the buccal dataset contained substantially more samples than the cfDNA dataset (1,187 versus 100), direct quantitative comparisons between datasets should be interpreted with this imbalance in mind. Nevertheless, distinct nucleotide enrichment patterns were observed across correlation, motif, and predictive analyses in both sample types. Across both datasets, the major k-mer enrichment patterns were largely preserved following truncation, suggesting that the observed sequence signatures were not predominantly driven by read-start composition. GC-enrichment and AT-enrichment patterns associated with age-correlated k-mers were further supported by WebLogo and MEME motif analyses. Similar trends were reproduced in synthetic datasets generated under controlled compositional assumptions. Collectively, these findings support the presence of reproducible age-associated sequence composition signatures that differ between canine buccal and cfDNA whole-genome sequencing datasets.

## 4. Discussion

Analysis of canine whole-genome sequencing datasets revealed age-associated signals in both buccal DNA and cfDNA, with substantially stronger patterns observed in buccal samples than in cfDNA. These findings complement previous studies demonstrating that age-associated molecular information can be captured through DNA methylation-based biomarkers [3, 4] and cfDNA-based aging biomarkers [8]. While most existing aging biomarkers rely on targeted molecular measurements, these findings suggest that age-related signals may be detectable through alignment-free analysis of sequence composition. The stronger patterns observed in buccal DNA further suggest that the effectiveness of k-mer-based approaches is influenced by the biological origin and composition of the sampled DNA. The differences between sample types likely reflect distinct biological processes, such as microbial community remodeling in buccal samples and chromatin reorganization in cfDNA, each of which leaves characteristic signatures in sequence composition.

Previous studies have demonstrated age-associated changes in canine oral microbiome composition [17, 18]. Aging has been associated with alterations in immune function, chronic inflammation, oral physiology, and host-microbe interactions that can influence the abundance of specific microbial taxa [13–15]. These changes likely contribute to the restructuring of the oral communities observed with age. Since microbial genomes vary substantially in nucleotide composition, including GC content, age-associated shifts in microbial community structure may contribute to the GC-enriched and AT-enriched k-mer patterns observed in buccal DNA. The successful reproduction of these patterns using simulations incorporating shifts in GC-rich bacterial sequences further supports the hypothesis that age-associated changes in microbial community composition contribute to the observed nucleotide enrichment patterns.

Age-associated changes in cfDNA likely reflect underlying alterations in chromatin organization. Since DNA wrapped around nucleosomes is protected from degradation, cfDNA fragmentation patterns retain indirect information about chromatin structure at the time of cell death [9]. With age, progressive heterochromatin loss and retrotransposon derepression alter chromatin organization across specific genomic regions, reducing nucleosome occupancy in regions that were previously tightly packed and shifting the population of DNA fragments released into circulation [1, 16]. This effectively means that cfDNA fragmentation patterns function as a liquid biopsy of age-related chromatin remodeling and the aging chromatin landscape directly determines which DNA is protected from degradation as well as which appears in circulation. Studies have shown that these structural shifts are detectable in cfDNA coverage patterns, including altered nucleosome positioning around transcription start and termination sites and increased accessibility of heterochromatic compartments [1, 8, 19]. Nevertheless, the heterogeneous tissue origins of cfDNA are likely to dilute these signals relative to single-tissue sources such as buccal swabs, which may explain the weaker age-associated patterns and reduced predictive performance observed in machine learning.

The age-associated signals observed in canine buccal samples may also have implications for the interpretation of analogous studies in humans. Dogs commonly experience age-associated changes in oral health and oral microbial community composition, which may contribute substantially to the sequence composition patterns detected in buccal DNA [17, 18]. In contrast, human populations may exhibit more consistent oral hygiene practices and dental care, potentially reducing the magnitude of age-associated microbiome shifts and their contribution to buccal sequence composition patterns. Consequently, the contribution of microbial communities to buccal aging signals may differ between species, and k-mer-based approaches applied to human buccal DNA may capture predominantly host cellular aging signals rather than microbiome-driven compositional changes. Future comparative studies across canine and human cohorts will be important for determining whether the strong buccal signals observed here represent a general feature of aging or reflect species-specific changes in oral microbial ecology.

Motif and sequence composition analyses further support these observations, as buccal-derived age-associated k-mers exhibit stronger motif conservation and more pronounced nucleotide enrichment pattern than cfDNA-derived k-mers. Notably, age-associated nucleotide composition trends differed between sample types, with positively correlated buccal k-mers exhibiting GC enrichment and negatively correlated buccal k-mers exhibiting AT enrichment, whereas the opposite pattern was observed in cfDNA-derived k-mers. These contrasting enrichment profiles suggest that distinct biological processes may underlie age-associated sequence composition changes across the two sample types. Furthermore, synthetic datasets generated under biologically motivated compositional assumptions successfully reproduced major nucleotide enrichment trends observed in empirical datasets. Although these simulations do not establish causality, they demonstrate that simple changes in underlying sequence compositions are sufficient to generate patterns resembling those observed in real sequencing data.

To assess whether age-associated signals are retained within read-start sequence composition, truncated data used only the first seven bases of each read. Comparison of truncated and raw datasets revealed minimal differences in correlation patterns, motif compositions, and age-prediction performance. These findings suggest that age-associated sequence signals are robust to preprocessing and are largely retained within read-start sequence composition, while remaining substantially stronger in buccal DNA than in cfDNA. This has potential implications for computational efficiency, as read-start k-mer analysis requires substantially less sequence information than full-length read analysis while preserving the majority of age-associated signals.

Several limitations should be considered when interpreting these findings. First, the cfDNA cohort size consisted only of 100 samples, whereas the buccal cohort had 1,187 samples. This substantial difference in sample size likely affected statistical power and may have contributed to observed disparities in significant k-mer counts and predictive performance between datasets. Second, all analyses were performed using canine samples, and therefore the generalizability of these findings to other species remains uncertain. Third, the two cohorts differed in age distribution. The buccal cohort was enriched for younger dogs, with a median age of 1.5 years and most samples falling below five years, whereas the cfDNA cohort was concentrated in older dogs, with a median age of 10 years. Consequently, age-associated k-mer signals may reflect different phases of aging trajectory across datasets, and the observed differences in signal strength may partly reflect this distributional disparity rather than intrinsic properties of sample types alone. Fourth, adapter removal was not performed prior to k-mer analysis, potentially introducing adapter-derived sequences into the raw-read datasets. Although this is less likely to affect the truncated datasets, which retained only the first seven nucleotides of each read, adapter contamination may have influenced the k-mer frequency distributions and age-associated correlations observed in full-length reads. Accordingly, comparisons between raw and truncated datasets should be interpreted with consideration of these preprocessing differences. Finally, the observed relationships are correlative and do not identify the molecular processes responsible for the detected sequence patterns. Future mechanistic studies integrating complementary molecular measurements may help determine the biological origins of the observed age-associated k-mer signals.

Future studies should first evaluate larger and more diverse cfDNA cohorts to improve statistical power and validate robustness of identified signals. Incorporating canine cellular blood DNA would help distinguish the effects of DNA fragmentation from tissue-specific sequence composition patterns. These comparisons could then be combined with complementary datasets, including DNA methylation, fragmentomic, transcriptomic, and microbiome profiling data, to investigate the biological mechanisms underlying age-associated k-mer signals. Ultimately, evaluating these patterns across additional species will help determine whether the observed sequence-level aging signals represent conserved features of the aging process.

## 5. Supplementary

Supplementary Figures 1 and Figure 2 are provided in the accompanying supplementary file.

