## Supplemental Data 1 for "Alignment-Free k-mer Analysis Reveals Age-Associated k-mer Signals in Canine Buccal DNA and Cell-Free DNA"

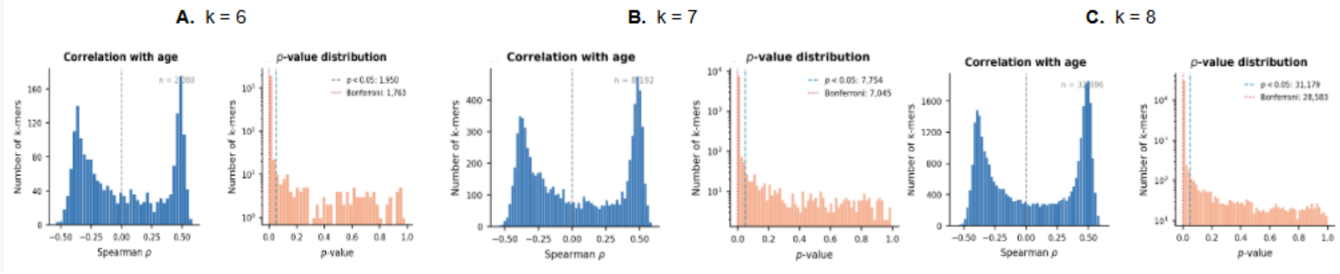


**Supplementary Figure 1.** Spearman correlation and p-value distributions of age-associated k-mers in raw buccal samples across k-mer sizes of 6, 7, and 8

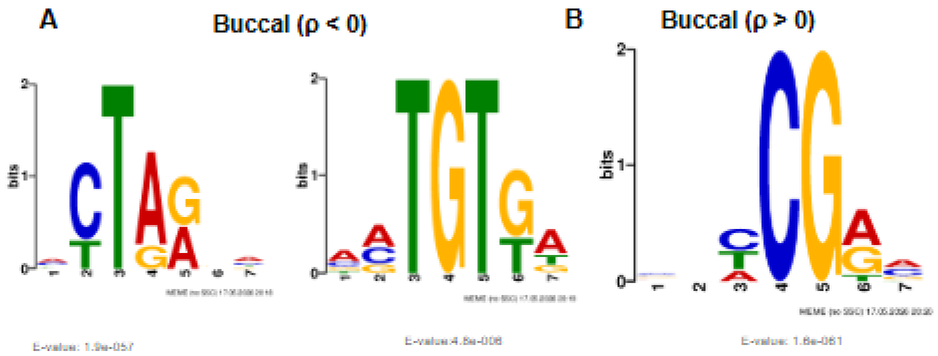


**Supplementary Figure 2.** Motif discovery in age-associated raw k-mers using MEME
